# RNAtranslator: Modeling protein-conditional RNA design as sequence-to-sequence natural language translation

**DOI:** 10.1101/2025.03.04.641375

**Authors:** Sobhan Shukueian Tabrizi, Sina Barazandeh, Helyasadat Hashemi Aghdam, A. Ercüment Çiçek

## Abstract

Protein-RNA interactions are essential in gene regulation, splicing, RNA stability, and translation, making RNA a promising therapeutic agent for targeting proteins, including those considered undruggable. However, designing RNA sequences that selectively bind to proteins remains a significant challenge due to the vast sequence space and limitations of current experimental and computational methods. Traditional approaches rely on in vitro selection techniques or computational models that require post-generation optimization, restricting their applicability to well-characterized proteins.

We introduce RNAtranslator, a generative language model that formulates protein-conditional RNA design as a sequence-to-sequence natural language translation problem for the first time. By learning a joint representation of RNA and protein interactions from large-scale datasets, RNAtranslator directly generates binding RNA sequences for any given protein target without the need for additional optimization. Our results demonstrate that RNAtranslator produces RNA sequences with natural-like properties, high novelty, and enhanced binding affinity compared to existing methods. This approach enables efficient RNA design for a wide range of proteins, paving the way for new RNA-based therapeutics and synthetic biology applications. The model and the code is released at github.com/ciceklab/RNAtranslator.

## 1 Introduction

Protein-RNA binding is an important component of the interactome and plays crucial roles in gene expression regulation [1], splicing [2], RNA stability [3], molecular degradation [4], RNA transport [5], and translation [6]. RNAs bind to proteins through either (i) RNA-binding domains (RBDs), which have rigid and well-defined 3D structures and selectively bind to specific RNA motifs, or (ii) Intrinsically Disordered Regions (IDRs), which have highly flexible structures, allowing them to interact dynamically with multiple binding RNA partners. The latter provides exciting opportunities for the design of novel RNA molecules that target and bind with proteins [7]. These can be used to inhibit, enhance, or modify RNA-protein interactions to restore functionality or eliminate disease-causing factors. RNA is a new and promising therapeutic agent to target proteins because only 15% of human proteins have binding pockets for ligands (small molecules) which are used as the common drug mechanism today [8]. This means that RNA has the potential to target proteins that have previously been deemed undruggable either with direct binding or by inhibiting the precursor RNAs. RNA-based drugs are also very favorable because (i) they have a lower immune system activation and inflammatory response with nucleoside-modifications; (ii) they have no genotoxicity risk as they do not integrate into the genome; (iii) they are relatively swift and inexpensive to produce with optimized chemistry and delivery [9].

The bottleneck in protein-conditional RNA design is the sheer number of possibilities for the candidate sequence, and unfortunately, current practice in the laboratory relies on manual and experimental design. The first strategy is to generate a large pool of candidate binding RNA sequences for a target using tools such as Primer-BLAST [10] followed by in vitro filtering of the library using experimental validation approaches such as SELEX (Systematic Evolution of Ligands by EXponential Enrichment) [11] which becomes more stringent in every validation cycle. The key element for success when it comes to selecting a good library to start with is based on luck, which is clearly not desirable. The second approach begins with an existing solution to a related problem, if one can be found, which is then optimized through the same in vitro test cycle. Both methods cost significant time and money and produce a suboptimal design. Computational techniques can aid design but due to their key limitations they are limited to proof-of-concept.

Most computational RNA design methods focus on *unconditional* RNA design. While early efforts solve the inverse folding problem via optimization [12, 13, 14, 15, 16, 17, 18, 19, 20], recent works use deep/reinforcement learning to design RNA structure (2D or 3D) [21, 22, 23, 24, 25, 26, 27] or RNA sequence [28, 29, 30, 31, 32]. Although the these methods can design synthetic RNA with natural-like properties such as GC content and minimum-free energy, only a few are capable of designing RNAs to target specific proteins (i.e., protein-conditional design). RNAFlow [24] is the only model that can design a *structure* to target a protein which uses a conditional flow-matching method. However, state-of-the-art structure designing models including RNAFlow cannot design novel molecules with variable length and the models can only recapitulate structures in the dataset [26]. This is because RNA is structurally very flexible compared to proteins and the paucity of ground truth structures bar training deep learning models [33]. Thus, the performances are random when designing conformationally diverse RNAs [24].

Protein-conditional RNA *sequence* design models do not suffer from training data scarcity and learn a latent distribution over widely available RNA sequence databases using architectures like generative adversarial networks [28, 30] and GPT-based language models [31]. However, conditioning on the target requires a post-generation optimization step with a binding affinity prediction method for the target protein. This means the current state-of-the-art procedure have key bottlenecks: (1) The two-step approach (generation+optimization) does not harness the full potential of learning better embeddings for RNA in relation to other molecules (targets); (2) The pipeline is dependent on the availability of RNA interaction data for the target and/or third-party tools that predict the binding affinity of the designed RNA with the target protein. For example, such Deep-Bind [14] models are available for only 732 proteins and DeepCLIP models [34] are available for 221 proteins. This means it is not possible to design RNA to target under-studied proteins or novel synthetic proteins. This is a crucial bottleneck for the applicability of this system as the number of proteins in the human body is estimated to be between 20,000 to several millions depending on the definition [35, 36, 37].

In this study, we propose a novel approach to overcome the challenges mentioned above. This first-of-its-kind method enables the design of binding RNA for *any* target protein. That is, for the first time, we formulate the problem of protein-conditional RNA design as a sequence-to-sequence *natural language translation* problem (not to be confused with biological process of protein generation). We use large-scale RNA-protein interaction datasets to train a generative language model, RNAtranslator, end-to-end. Like ChatGPT inputing a sentence in English and translating it into French, RNAtranslator inputs a target protein sequence and *translates* it to a novel binding RNA sequence. The model learns a latent space for both the RNA and the protein along with their interactions and thus, does not require any postprocessing or optimization method to tailor the design for binding to the target. Thus, the model is capable of designing RNA to bind with any target protein sequence. **For the first time, we enable designing RNA to target understudied, novel or synthetic proteins**.

Our results on targeting 10 proteins show that RNAtranslator is capable of generating RNAs which resemble natural binding RNAs with respect to length, GC content, minimum free energy (MFE) and ensemble free energy distributions, yet we also find that the designed sequences are novel. We observe that RNAtranslator is substantially better compared to the state-of-the-art methods and is capable of designing for any target protein. While stability is often inversely correlated with binding affinity, we also find that the designed RNAs have higher binding affinity to the target compared to the state-of-the-art using (i) binding affinity prediction, (i) docking prediction, and (iii) molecular dynamics simulation methods. We design RNAs to target proteins which (i) do not have any known RNA interactions (PRP4K) and (ii) the model has not observed in its training procedure (PRPF8) and observe that the designs have high binding affinity to their respective targets. This shows that the model has learned the sequence characteristics driving an interaction and has not overfit to known proteins. Designing for these cases is possible for the first time with RNAtranslator. We think the potential of RNAtranslator goes beyond this study and it will pave the way for modeling and designing for many other interactions and will be the first milestone in the new field of *molecular translation*. The model and the code is released at github.com/ciceklab/RNAtranslator.

## 2 Results

### 2.1 RNAtranslator Overview

RNAtranslator is an encoder-decoder transformer-based language model designed for RNA sequence generation conditioned on a target protein sequence input. The model aims to generate RNA sequences that exhibit strong binding affinity to target protein while maintaining structural stability and natural RNA-like properties. The architecture of RNAtranslator is shown in Figure 1. We trained the model in two steps: (i) we first trained it using 26 million RNA-protein interactions from the RNAInter dataset, which includes both computational and experimental data, and (ii) we then fine-tuned it with 12 million experimentally validated interactions. The model encoder is input with the target protein sequence. The decoder inputs the binding RNA sequence and the embedded protein sequence and learns to regenerate the RNA sequence from left to right. During inference, the model inputs the target protein sequence and generates novel RNA sequences that are likely to bind to it. We select the candidate sequence after sampling from this step.

**Figure 1.**
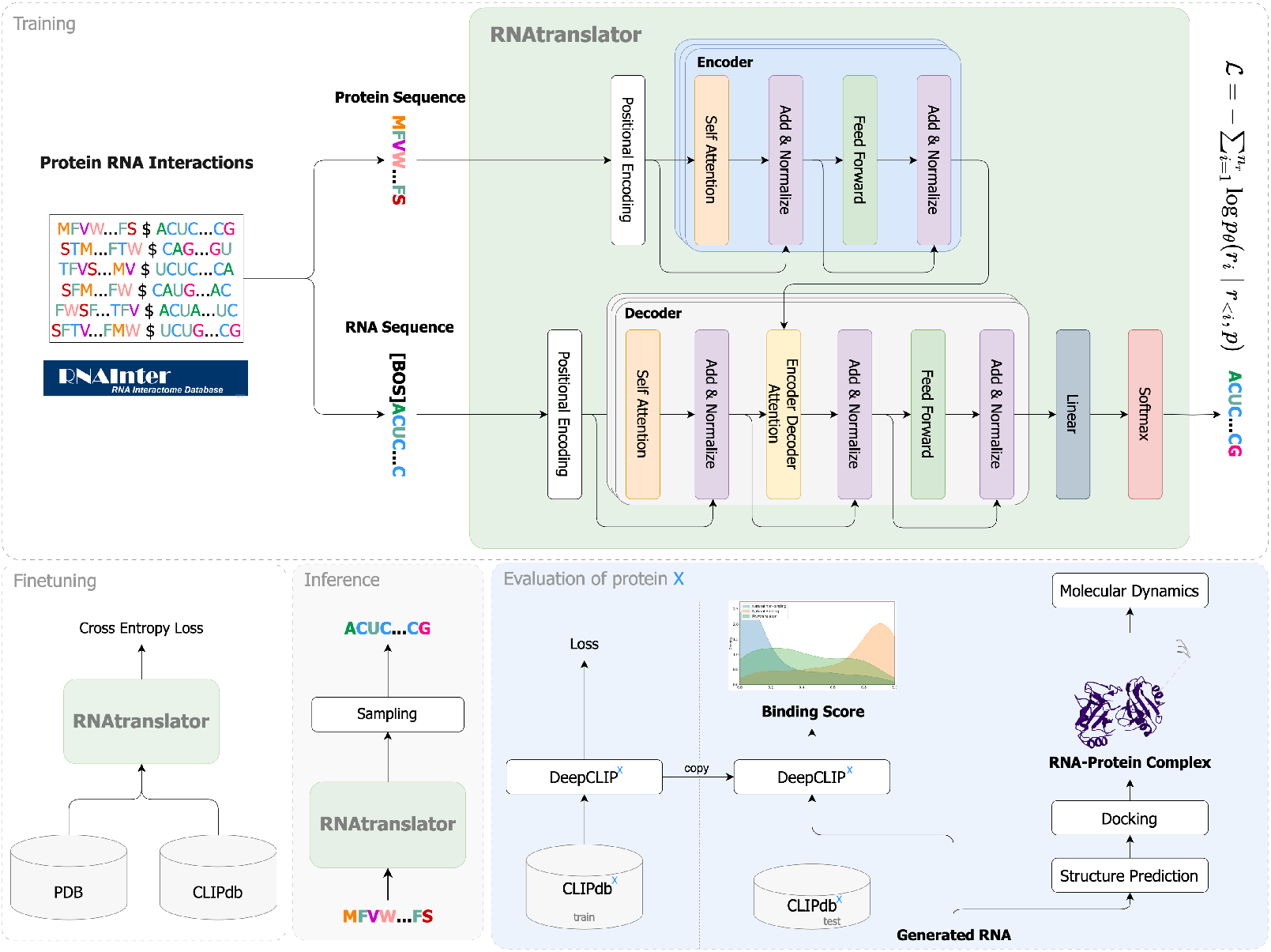
Overview of RNAtranslator pipeline: The model follows an encoder-decoder transformer framework, where the encoder processes the protein sequence using positional encoding and encoder blocks to extract an embedding. The decoder takes the RNA sequence input, applying selfattention, encoder-decoder attention, and feed-forward layers to learn to generate target-specific RNA sequences. Training occurs in two steps: Large scale pretraining with experimental and computationally predicted interactions followed by fine-tuning with experimental interactions. During inference, the model requires only the protein sequence as input and generates novel RNA sequences through iterative sampling. We evaluate designed RNAs based on predicted in-silico binding affinity to target protein X using DeepCLIP (trained on the target’s CLIPdb data), as well as by docking predictions between the RNA structure and target protein X structure. DeepCLIP^*X*^ indicates the model trained to predict binding affinity to X and CLIPdb^*X*^ indicates the subset of the dataset which includes only iteractions for protein X.

To evaluate our designs, we use in silico binding affinity predictors such as DeepCLIP [34] and docking tools to simulate the molecular interaction between the design and the target structure. We also use OpenMM [38] to simulate the dynamics of our design bound to the target protein. Molecular dynamics simulations help reveal how these complexes behave in a physiologically relevant environment rather than a static crystal structure. These simulations provide insights into conformational changes, hydrogen-bonding networks, and the thermodynamics of binding at atomic resolution by using force fields optimized for both proteins and RNA. Please see Section 4.6 for the details on the simulation.

In the upcoming subsections, we first evaluate the novelty of the RNA sequences generated using RNAtranslator. This step ensures that our generated RNAs are unique compared to existing known RNA sequences that bind with the target. Subsequently, we design RNAs to bind with RNA-binding proteins RBM5 and ELAVL1 proteins and compare their binding affinities with (i) RNA sequences generated by the state-of-the-art methods, (ii) natural known binders of these proteins, and (iii) randomly selected natural RNAs that are unlikely to bind. To further test the generalizability of RNAtranslator, we analyze its ability to design binding RNAs for unseen proteins. The PRPF8 and PRP4K interactions were not included in our training data set. In fact, PRP4K does not have any known RNA interaction set in the datasets. Thus, it would not be possible for the state-of-the-art methods to design an RNA for this protein. By predicting and validating RNA-protein interactions through docking and molecular dynamics simulations, we confirm that our approach can effectively target understudied and novel (e.g., synthetic) proteins. We also extend our analysis to 9 more RNA-binding proteins to determine the broader applicability of our RNA design method. Finally, we analyze the stability of our generated RNAs to confirm their structural robustness and thermodynamic favorability.

### 2.2 RNAtranslator generates novel and binding RNAs

We evaluate the novelty of generated RNA sequences by comparing them with natural binding RNAs using BLAST (Basic Local Alignment Search Tool). To optimize our analysis, we first separate the RNA sequences into short and long groups based on a 50-nucleotide threshold and adjust the BLAST word size—using 4 for short sequences and 7 for long ones. Next, we compare each sequence with the target proteins’ binding RNA database by applying the following thresholds: (i) 70% identity, (ii) a minimum alignment length of 15 base pairs, (iii) 50% query coverage, and (iv) an E-value of 0.1. Sequences that meet these criteria are classified as *similar to known*, while those that do not are considered novel.

Here, we compare RNAtranslator with only two other methods that can perform protein-conditional RNA design using the target sequence: RNAGEN [28] and GenerRNA [31]. RNA-GEN is a generative adversarial neural network that generates RNA sequences *de novo*. In a post-processing step seed RNAs are then optimized to bind to the target using a binding affinity predictor’s rewards. GenerRNA is a GPT-based generative language model which can design RNAs *de novo* as well. The model needs to be fine-tuned with the target protein’s RNA interactions for conditional design.

We generate 500 RNAs using each method and compare the percentage of novel RNA sequences in the designed sets using the criteria mentioned above. We select two proteins with substantially different numbers of known RNA interactions from the CLIPdb database: RBM5, which has a relatively small set of interactions (3, 917 interactions), and ELAVL1, with a notably large set of interactions (1, 079, 145 interactions). This selection allows us to investigate how the size of the training dataset affects the novelty of generated RNA sequences. RBM5 protein is an RNA binding protein that plays a role in apoptosis through pre-mRNA splicing of target genes. Its role is well characterized in cancer and is a candidate drug target for other conditions such as osteoprosis [39] and central nervous system disorders [40]. ELAVL1 is also an RNA binding protein that stabilizes mRNAs by binding to AU-rich regions. It has also been associated with cancer [41]. Thus, successfully designing RNA to target these proteins holds therapeutical potential.

Our results show that 83.4% of the RNAtranslator-designed RNAs for RBM5 are novel. For ELAVL1 71.2% of the designed sequences are novel. GenerRNA yields a substantially lower novelty rate, suggesting the fine-tuning procedure for the target protein leading to overfitting. Only 56.76% and 56.4% of the designed sequences are novel for RBM5 and ELAVL1, respectively. RNA-GEN achieves a comparable novelty rate for RBM5 (81.64%) and higher novelty rate for ELAVL1 (85.94%). Note that RNAGEN does not directly use interaction data but uses a third-party affinity predictor tool to customize the designed RNA which makes it less likely to produce known RNAs.

We filtered out known RNAs from all groups and compared the binding affinities of novel RNA sequences to natural RNAs using DeepCLIP. For each protein target, we selected the RNA sequence with the highest binding affinity score and performed molecular docking analysis using HDOCKlite, which provided the top 100 docking models indicating potential binding positions. In Figure 2, panels (a) and (c) show the distributions of binding affinities for RBM5 and ELAVL1 proteins, respectively. Overall, all methods performed better for ELAVL1, likely due to significantly larger training data. Additionally, RNAs generated by RNAtranslator generally showed higher binding affinity than RNAs produced by other methods. Panels (b) and (d) show docking score distributions for the top 100 docking models of RBM5 and ELAVL1, respectively, where lower scores indicate better binding. For RBM5, while other tools’ docking score distributions resemble natural binders’ distribution, RNAtranslator-generated RNAs achieve a distribution whose mean is lower than even experimentally validated natural RNAs’ distribution. This highlights the effectiveness of our model in designieng high-binding-affinity-RNAs with a relatively small set of RNA interactions. For ELAVL1, our model achieves the closest mean binding affinity to the natural binders of the protein.

**Figure 2.**
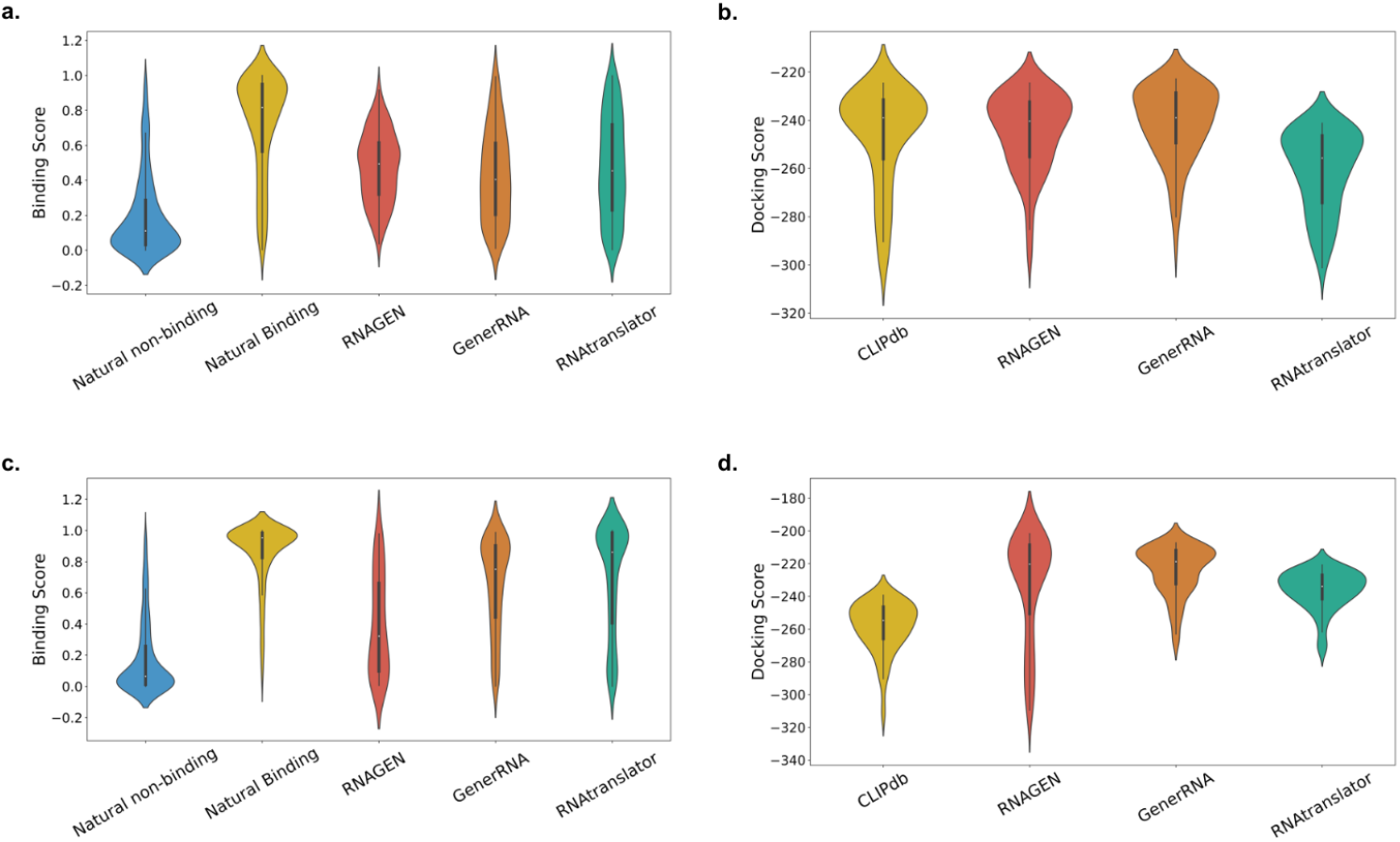
Comparison of predicted binding affinities between generated and natural RNAs for RBM5 and ELAVL1 proteins as targets. (a) Distribution of binding affinities for RBM5 protein predicted by DeepCLIP. (b) HDOCKlite scores for the top 100 models for RBM5-RNA complexes. (c) Distribution of binding affinities for ELAVL1 protein predicted by DeepCLIP. (d) HDOCKlite scores for the top 100 models of ELAVL1-RNA complexes.

We also evaluate how the proteins RBM5 and ELAVL1 interact with RNAs designed by RNA-translator using molecular dynamics simulations (see Section 4.6). As shown in Figure 3a, RNAs from RNAtranslator form more contacts and have lower (better) binding energies with both RBM5 and ELAVL1 compared to natural binding RNAs and random natural RNAs. We also analyze how stable these protein-RNA complexes are during the simulation in Figure 3b using FEL plots which show the unbound ELAVL1 protein has a maximum FEL depth of 14.96*kcal/mol*, whereas binding to RNAtranslator yields a deeper well at 15.97*kcal/mol*, and the natural binding RNA reaches 15.82*kcal/mol*. Since deeper wells indicate greater stability, both RNAs stabilize ELAVL1 compared to its unbound form. Interestingly, the FEL analysis for RBM5 reveals the opposite trend. The unbound RBM5 protein has the deepest well at 15.45*kcal/mol*, compared to 15.08*kcal/mol* for the RNAtranslator-bound complex and 15.14*kcal/mol* for the natural binding RNA complex. Thus, RBM5 appears intrinsically more stable when unbound. Nonetheless, among the RNA-bound complexes, RNAtranslator still produces a slightly deeper (more stable) energy basin than the natural binding RNA, consistent with its stronger binding energy profile and higher number of molecular contacts.

**Figure 3.**
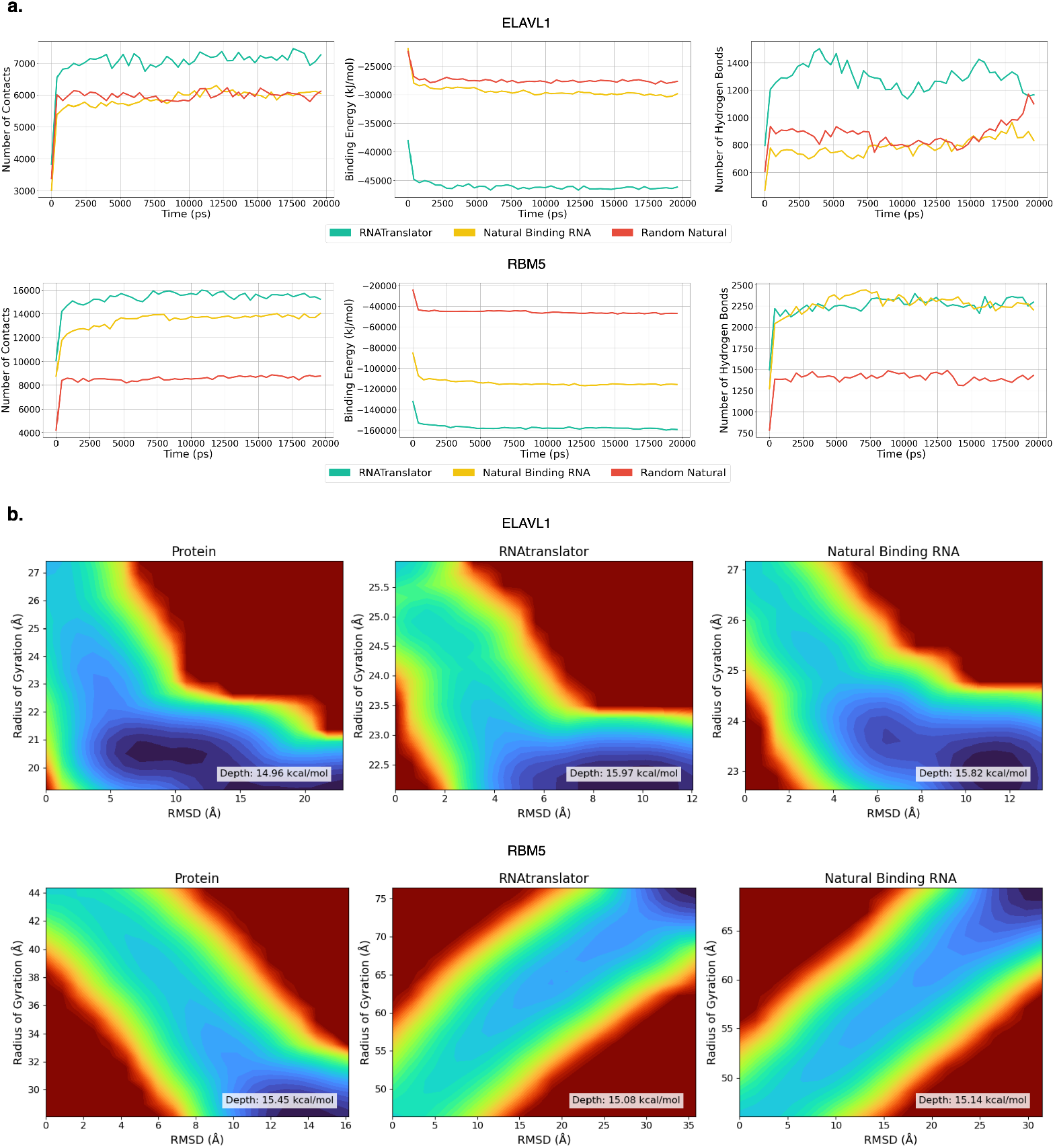
Molecular dynamics simulations showing interactions of RBM5 and ELAVL1 with RNAs generated by RNAtranslator and Natural RNAs: Panels (a) shows that RNAs designed by RNAtranslator form more contacts and have lower (better) binding energies compared to natural and random natural RNAs. Panel (b) shows the stability analysis; ELAVL1 complexes are more stable (deeper regions in the plot), while RBM5 complexes show reduced stability with larger structural changes.

While we show that RNAtranslator has novel RNAs with better binding affinity, it would have been just another method with incremental improvements. The most important advantage and novelty of the our method over the others is that it does not require additional information for the target protein. This enables RNAtranslator to design RNAs to target understudied or novel/synthetic proteins. Note that to use RNAGEN and GenerRNA, we need post-processing of the model or the designed RNAs. That is, we need to optimize the RNAs generated by RNAGEN to have them bind to RBM5 and ELAVL1 using a specific protein-target specific binding affinity predictor trained. Similarly, we need to fine-tune the base GenerRNA model with RNAs that are known to bind to RBM5 and ELAVL1, respectively. These tools/data might not be available at all, prohibiting the conditional design. RNAtranslator is free from these steps, which enables it to generalize to any protein sequence and to work in the absence of this information. We demonstrate this in the next section with a protein that does not have a binding affinity predictor or known RNA interactions. Thus, RNAGEN and GenerRNA cannot design RNAs to target this protein.

### 2.3 RNAtranslator can design for a protein without available RNA interaction data

To test how well our model generalizes, we evaluated it on two proteins not included in the training: PRPF8 and PRP4K. PRP4K is a kinase responsible for pre-mRNA splicing, signal transduction, and tumor suppression [42]. There are no known RNA interactions for this protein in RNA interaction databases which means it is not possible to design RNA to target PRP4K with current methods. PRPF8, on the other hand, is a core component of the U5 small nuclear ribonucleoprotein (snRNP) complex within the spliceosome. It is essential for the precise removal of introns during pre-mRNA splicing by orchestrating the interactions between pre-mRNAs and small nuclear RNAs (snRNAs) [43]. There are 27, 046 interactions for this protein in the CLIPdb database, which we did not use during training but instead used to evaluate our model with the DeepCLIP method

We cannot predict the binding affinity of the designed RNA sequences for PRP4K using DeepCLIP as there is no interaction data to train such an affinity predictor. Thus, we analyze the binding affinity using (i) RNA-protein docking methods which computationally model the complex using structures of the interactors, and (ii) performing molecular dynamics (MD) simulations of the predicted complex obtained from the docking tool. The pipleline of evaluating generated RNA for this protein is shown in Figure 4a, first we obtain the ground-truth crystal structure of PRP4K available in the Protein Data Bank (PDB entry: 4IAN, determined by X-ray crystallography at 2.44 Å resolution), then we predict the structure using RhoFold+ [44]. Using HDOCKlite [45], we model the RNA-protein complex. Finally, we perform MD simulations using OpenMM [46]. Analysis of hydrogen-bond and contact plots in Figure 4b reveals that the RNAtranslator bound PRP4K consistently forms more contacts compared to randomly selected natural RNAs. Throughout the simulation, the RNAtranslator-generated RNA maintains more hydrogen bonds relative to these other RNAs. This increased number of hydrogen bonds indicates stronger interactions, which is further confirmed by the binding energy analysis. The RNAtranslator bound PRP4K demonstrates binding energies that are more negative than the natural RNAs, highlighting its better binding affinity and stability. Additionally, the free-energy landscape (FEL) analysis shown in Figure 4c supports these findings. The FEL analysis compares the depth of energy basins for both bound and unbound states. The unbound protein shows a maximum basin depth of approximately 15.72*kcal/mol*, whereas the RNAtranslator-generated-RNA-protein complex achieves a notably deeper basin of around 16.59*kcal/mo*l which is the deepest observed among all tested complexes. In comparison, the random-natural RNA complexes reach depths of approximately 16.20, 15.82, and 15.90*kcal/mol*. These depths are deeper than the unbound protein state but still shallower than the protein of RNAtranslator-generated-RNA-protein complex.

**Figure 4.**
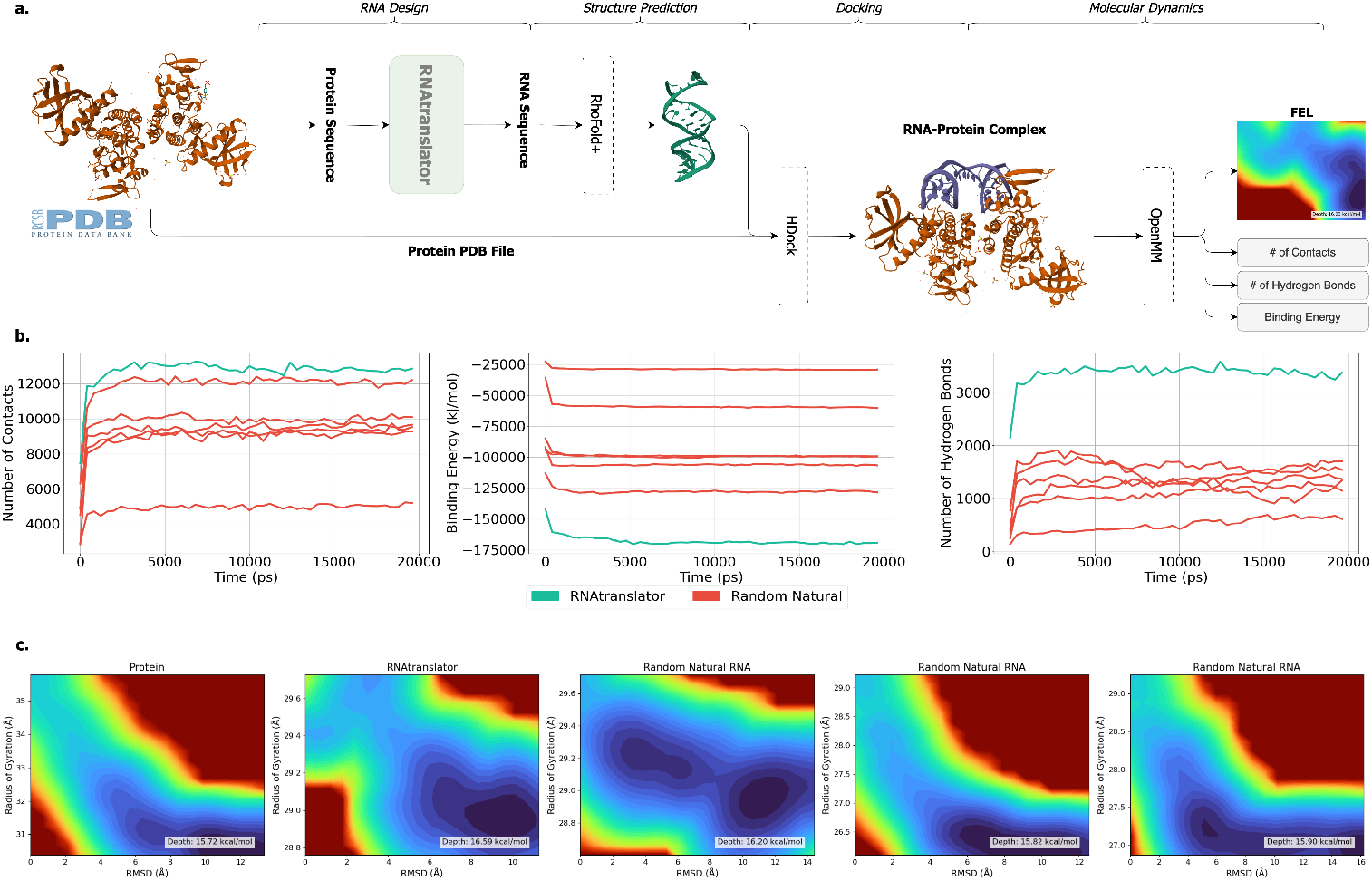
Comparison of RNA-protein interactions for PRP4K: (a) PRP4K bound RNA design and evaluation pipeline: first, we generate RNA for the ground-truth crystal structure of PRP4K (4IAN), predict the structure using RhoFold+, model the RNA-protein complex using HDOCKlite, and finally perform MD simulations using OpenMM. (b) MD simulation analysis demonstrates that RNAs generated by RNAtranslator maintain more contacts and hydrogen bonds and have lower (stronger) binding energies compared to six randomly selected natural RNAs, indicating enhanced binding affinity. (c) Free-energy landscape (FEL) plots comparing unbound and RNA-bound states of the protein, where the RNAtranslator–protein complex achieves the deepest basin (approximately 16.59*kcal/mol*) among all tested complexes.

PRPF8 is a large protein containing 2335 amino acids, to fit to our model we truncate the first 400 amino acids, which are not annotated as binding domains in UniProt, reducing the protein to 1935 amino acids. Using RNAtranslator, we generated 500 RNAs for PRPF8 and compared them with natural binding RNAs using the DeepCLIP model. Figure 5a shows that the distribution of binding scores for RNAtranslator RNAs closely matches that of natural binding RNAs. Figure 5b illustrates the predicted best complex formed between PRPF8 and a RNAtranslator RNA, the structure of PRPF8 is predicted using AlphaFold3 [47]. Additionally, through molecular dynamics simulations, we analyzed the interactions within the complexes. The results in Figure 5c show that, while the number of contacts is similar among RNAtranslator, natural binding, and non-binding RNAs, the binding energies of the RNAtranslator and natural binding RNAs are substantially lower, confirming their stronger binding interactions compared to non-binding RNAs. FELs in the Figure 5 (panel d), show that the unbound protein has a global minimum depth of 15.89*kcal/mol*. In contrast, the protein bound to RNAtranslator achieves a deeper minimum of 16.23*kcal/mol*, while the protein bound to the natural binding RNA reaches 16.03*kcal/mol*. These higher FEL depths indicate more stable conformations when the protein is bound to RNA.

**Figure 5.**
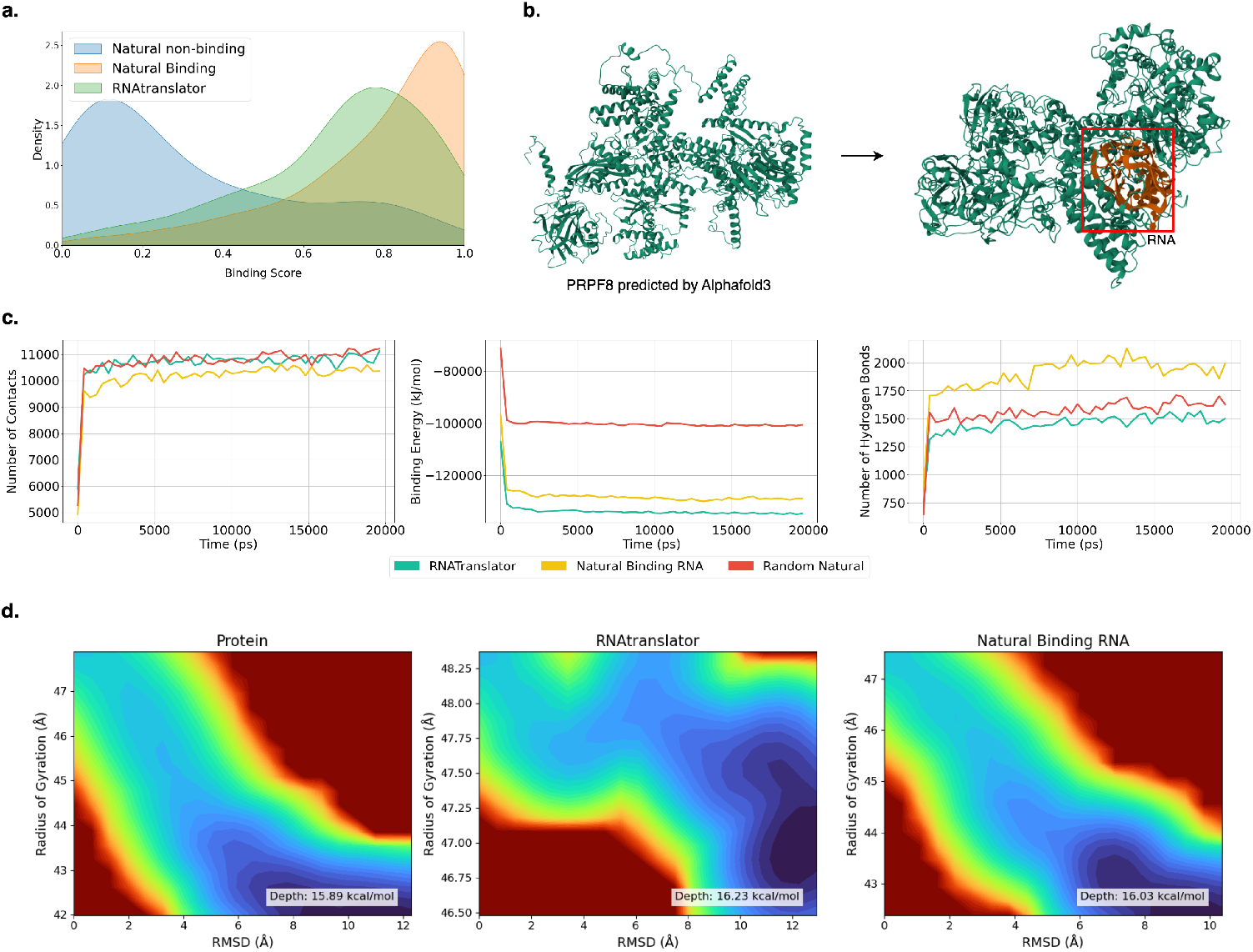
Analysis of PRPF8 interactions: (a) Distribution of binding scores showing RNAtranslator-generated RNAs closely resemble natural binding RNAs. (b) Predicted optimal PRPF8-RNA complex structure. (c) Molecular dynamics analysis reveals similar numbers of contacts but significantly lower binding energies for RNAtranslator and natural binding RNAs compared to non-binding RNAs, highlighting stronger interactions. (d)Free-energy landscape (FEL) plots show the RNAtranslator–protein complex achieves the deepest energy basin (approximately 16.23*kcal/mol*), closely followed by the natural binding RNA, both being deeper than the unbound protein, indicating increased stability.

### 2.4 RNAtranslator can design RNAs with high binding affinity to Nine targets

We evaluate the binding affinity of RNAs generated by RNAtranslator for nine proteins listed in CLIPdb. Specifically, we generate 128 RNAs each for ZC3H7B, AGO2, U2AF2, RBM5, HNRNPA1, TARDBP, MOV10, and SRSF1. By assessing these RNAs using the DeepCLIP model, we analyze the distribution of binding scores, allowing us to compare RNAtranslator-generated RNAs to natural binding and natural non-binding RNA sequences. These natural binding and non-binding RNAs are selected from an independent test set, ensuring that the DeepCLIP model has not been trained with these specific sequences, thus maintaining the integrity of our assessment.

As shown in Figure 6, RNAtranslator-generated RNAs generally show binding affinities comparable to naturally binding RNAs for most targets, notably for proteins such as AGO2, and SRSF1, where distributions overlap substantially. This overlap indicates that the sequences produced by RNAtranslator have a binding profile similar to naturally occurring RNAs, reflecting the model’s accuracy in capturing essential characteristics necessary for effective protein binding. Furthermore, a substantial difference between the distributions of natural binding and non-binding RNAs supports the robustness of the DeepCLIP predictions, confirming its effectiveness in distinguishing functionally relevant RNA-protein interactions. Together, these observations emphasize RNAtrans-lator’s capability to produce sequences with meaningful binding affinities across a range of protein targets, underscoring its utility for advancing research in RNA biology and developing RNA-based therapeutic strategies.

**Figure 6.**
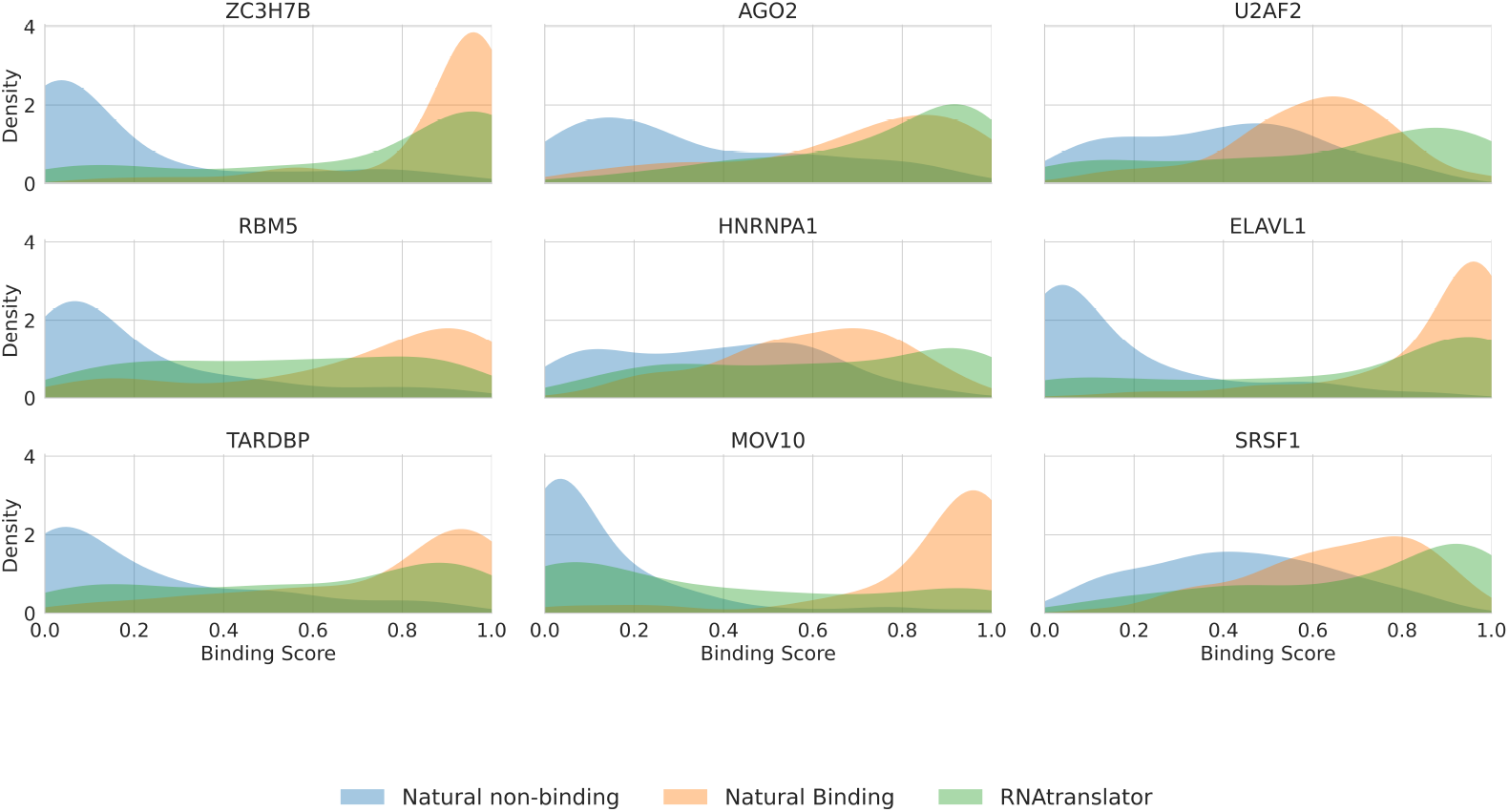
Binding affinity comparison of RNAtranslator-generated RNAs and natural RNAs. RNAtranslator-generated RNAs show high binding affinities across nine protein targets. The distributions of binding scores indicate that RNAtranslator-generated RNAs generally show affinities comparable to naturally binding RNAs, with substantial overlaps observed for most proteins.

### 2.5 RNAtranslator-designed RNAs are stable

To evaluate the stability of RNAtranslator-generated RNA sequences, we analyzed key stability metrics, including free energy, GC content, and ensemble free energy unfolding. The Minimum Free Energy (MFE) serves as an indicator of RNA stability by predicting the most thermodynamically favorable secondary structure. Lower MFE values suggest more stable conformations, as they represent the minimum energy required for an RNA molecule to fold [48]. Furthermore, GC content was examined as a complementary measure, since a higher GC content enhances structural robustness through stronger base-pairing and higher melting temperatures [49].

Our results show that RNAtranslator-generated RNAs are stable with respect to these metrics. The analysis of GC content in Figure 7 shows that natural binding RNAs tend to have a similar GC content compared to nonbinding natural RNAs, as expected. RNAs designed by RNAtranslator have even a stronger GC composition compared to natural binding RNAs in four columns. Our analysis on MFE also further supports these findings. As shown in Figure 7, RNAtranslator generated RNAs achieve a distribution slightly more stable than natural RNAs. This suggests that RNAs generated with RNAtranslator have a better energy landscape, which enhances their overall foldability and thermodynamic resilience. To provide a more comprehensive assessment of RNA stability, we incorporate the free energy unfolding of the ensemble (Δ*G*_*ensemble*_), which considers the entire thermodynamic ensemble of possible secondary conformations rather than just the lowest energy structure using ViennaRNA[50]. RNAtranslator generated RNAs have a distribution that tend to be slightly more negative than natural RNAs (Figure 7). The length of the RNA sequence also affects the secondary structure and binding affinity. It is also an important factor for diversity and novelty of the design. For this reason, we also compare the RNA length distributions across different groups. Again, as can be seen in Figure 7, RNAtranslator-generated RNAs span broader range than natural binding RNAs with higher median compared to natural non-binding RNAs. Stability analyses of RNAs generated for other five protein targets are presented in Supplementary Figure 3.

**Figure 7.**
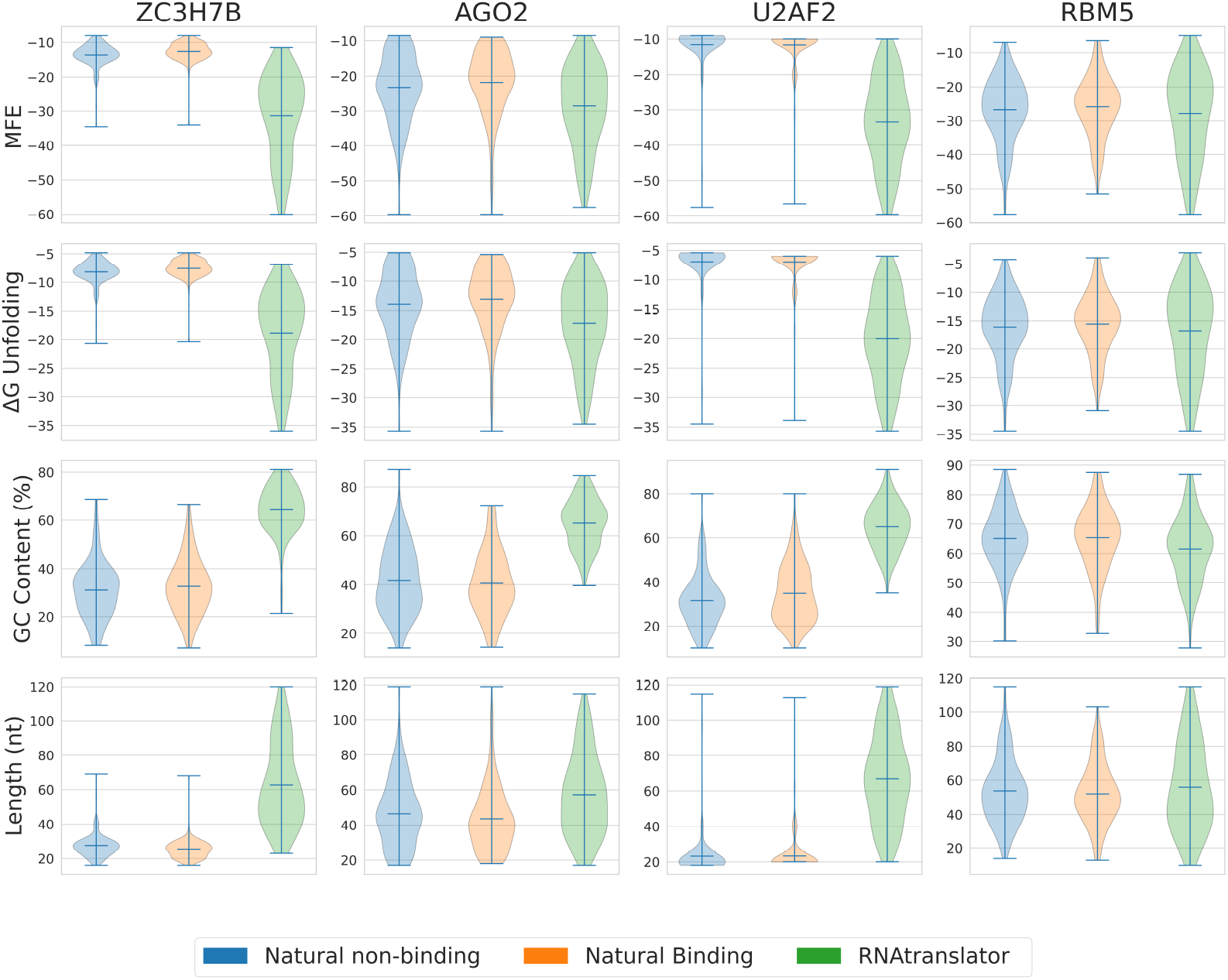
Stability analysis of RNAtranslator generated RNAs compared to natural RNAs. Minimum Free Energy (MFE) distributions is shown in the first row, where RNAtranslatorgenerated RNAs achieve stability levels comparable to natural binding RNAs, reinforcing their thermodynamic favorability. Ensemble Free Energy (Δ*G*_*ensemble*_) distributions, further validating the structural stability of RNAtranslator-generated RNAs in a thermodynamic ensemble. GC content comparison, indicating that RNAtranslator-generated RNAs closely match the natural binding RNAs, suggesting structural robustness. Distribution of RNA sequence lengths across different groups, showing that RNAtranslator-generated RNAs exhibit a broad yet biologically relevant length distribution.

Overall, compared to other methods, RNAtranslator generated RNAs display a more stable energy distribution with a large span of sequence lengths where RNAtranslator has a slight advantage. This reinforces RNAtranslator’s ability to adopt biologically relevant folding patterns under diverse conditions.

## 3 Discussion

We introduce a novel approach to protein-conditional RNA sequence design by formulating the problem as a natural language translation task. Unlike existing methods that rely on separate generation and optimization steps, our model, RNAtranslator, learns a unified latent space for RNA and protein interactions, enabling direct and target-specific RNA design. The results demonstrate that RNAtranslator outperforms state-of-the-art approaches in the generation of natural-like RNA sequences while improving binding affinity to the target. Our results suggest that the designed RNAs maintain stability while preserving sequence and length diversity. This balance is important, as excessive stability may lead to undesired rigidity which bars effective binding, while high flexiblity may lack specificity in the design. Additionally, our docking and binding affinity evaluations confirm that RNAtranslator designs natural-like RNA sequences with superior binding potential compared to existing models.

The advantage and the novelty of RNAtranslator lies in its ability to generalize beyond predefined RNA-protein interactions. Traditional RNA design methods rely on either (1) experimental selection processes, which are costly and time-intensive, or (2) computational models that suffer from data scarcity and the need for post-generation optimization. We eliminate these bottlenecks by using large-scale RNA-protein interaction datasets and end-to-end training on paired data, allowing the model to learn implicit binding rules without manual intervention. Moreover, RNA-translator does not rely on third-party binding affinity predictors, which are rarely available only for a few hundred well-studied proteins. For instance, we had to optimize the RNAs generated by RNAGEN using a binding affinity predictor trained specifically for RBM5 and we had to fine-tue the GenerRNA method with RNAs that are known to bind to RBM5. RNAtranslator does not require any of these steps as it was traine with large scale interaction data, end-to-end. This allows RNAtranslator to expand the scope of RNA-based therapeutics.

Despite these promising results, there are several areas for further improvement. One challenge is the scalability of training large language models on biological sequences, given the computational complexity of RNA-protein interactions. Future work could explore techniques such as parameter-efficient fine-tuning or self-supervised learning on larger, unlabeled datasets to further enhance performance. Another limitation is the lack of experimentally validated binding data for novel RNA designs. While our computational assessments suggest high-affinity interactions and molecular dynamics (MD) simulations can offer valuable insights into how molecules bind and how stable they are, they are not perfectly accurate. Errors may arise from choices such as force-field parameters, simulation length (which may not capture all relevant conformational changes), and the specific analysis methods used. Additionally, initial conditions, boundary effects, and the inherent limitations of classical physics approximations can introduce uncertainties. Nonetheless, MD results can still provide a reasonable, hypothesis-generating view of intermolecular interactions and stability trends, even if the absolute values should be treated with caution., experimental validation through techniques such as surface plasmon resonance or isothermal titration caliometry will be necessary to confirm binding efficacy in vitro. Despite these promising results, there are several areas for further improvement. One challenge is the scalability of training large language models on biological sequences, given the computational complexity of RNA-protein interactions. Future work could explore techniques such as parameter-efficient fine-tuning or self-supervised learning on larger, unlabeled datasets to further enhance performance. Another limitation is the lack of experimentally validated binding data for novel RNA designs. While our computational assessments suggest high-affinity interactions and molecular dynamics (MD) simulations offer insights into how molecules bind and how stable they are, they are not perfectly accurate. Also, errors may arise from initial conditions, boundary effects, and the inherent limitations of classical physics approximations that can introduce uncertainties. Experimental validation through techniques such as surface plasmon resonance or isothermal titration caliometry will be necessary to confirm binding efficacy in vitro [51].

In summary, our study presents a paradigm shift in protein-conditional RNA design by introducing an end-to-end generative approach that eliminates the need for post-optimization or fine-tuning. RNAtranslator successfully designs RNA sequences with enhanced stability, diversity, and binding affinity, overcoming key challenges in existing computational and experimental methods. Our findings suggest that this framework will play a pivotal role in the future of RNA therapeutics, molecular engineering, and synthetic biology.

## 4 Methods

### 4.1 Dataset

We collect RNA-Protein interaction data primarily from the RNAInter Database [52], which provides approximately 26 million interactions, combining both experimentally validated and computationally predicted RNA-protein interactions (RPIs). The dataset predominantly includes human interactions obtained from diverse high-throughput experimental methods, such as CLIP-seq, RIP-seq, and yeast three-hybrid assays, covering various RNA types like mRNA, lncRNA, miRNA, and circRNA.

To ensure data quality and reduce errors associated with computational predictions in RNAInter, we use high-quality interaction data from CLIPdb and the Protein Data Bank (PDB) complexes for finetuning RNAtranslator. Specifically, we obtain human interaction data from CLIPdb via the POTAR3 [53] repository. We initially deduplicate RNAs associated with each protein using CD-HIT-EST with a 90% identity threshold. After an analysis of the CLIPdb dataset, as shown in Supplementary Figure 1a there is a significant imbalance among proteins in terms of interaction counts, ranging from as few as 85 interactions (EZH2 protein) to over 1 million interactions (1, 079, 145 for ELAVL1 protein). As that such imbalance could bias our model, we use an oversampling strategy to achieve equal representation across all proteins. However, to prevent oversampling from disproportionately amplifying specific RNA sequences, we cluster the RNAs across the entire dataset and apply a weighted sampling method. This method prioritizes interactions involving less common RNA clusters, thereby preserving RNA sequence diversity and ensuring balanced protein representation. From this refined dataset shown in Supplementary Figure 1b, we randomly sample a total of 12 million interactions to fine-tune our model.

In addition, we further augment our fine-tuning dataset using the PRI30K [54] dataset, curated from protein-RNA complexes available in the Protein Data Bank (PDB). This dataset consists of protein-RNA complexes structured into pairwise interactions, carefully processed into a non-redundant collection containing approximately 30, 000 unique interactions. From this PRI30K dataset, we randomly sample interactions.

### 4.2 Problem Formulation

Let 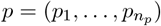 denote a protein sequence, where each *p*_*i*_ is a one or multiple amino acids token, and let *r* = (*r*_1_, …, *r*_*n*_*r*) be the corresponding RNA sequence we aim to generate. Our primary objective is to learn the conditional distribution *p*(*r*), which describes how likely an RNA sequence *r* is given a protein sequence *p*. By formulating this as a language translation problem, where the protein sequence is considered the source language and the RNA sequence is the target language, we used techniques from natural language processing (NLP) to model conditional protein-RNA design. We factorize *p*(*r*) via the chain rule:

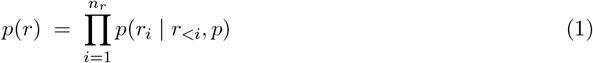

We then train a model with parameters *θ* that minimizes the negative log-likelihood (NLL) over a protein-RNA interaction dataset 𝒟 = {(*p*^(1)^, *r*^(1)^), …, (*p*^(|𝒟|)^, *r*^(|𝒟|)^)}.

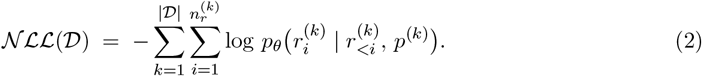

### 4.3 Model Architecture

We implement the RNAtranslator using the T5 architecture [55], a widely adopted encoder-decoder transformer designed for sequence-to-sequence tasks.

#### 4.3.1 Training

We feed the encoder with the protein sequence *p* to model the protein language and learn the information from a protein’s RNA-binding domains (RBDs). Before processing, each amino acid token is converted into a learned token embedding, using BPE tokenization to handle large protein vocabularies. Positional information is also encoded, enabling the model to differentiate between tokens in different sequence positions. Concretely, a tokenized protein sequence of length *n*_*p*_ is mapped to a sequence of embeddings, which is then passed through *l* layers of self-attention and feedforward blocks. The final output of the encoder is a matrix 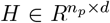 where *d* is the model dimension and encodes contextual information about the protein’s amino acid composition, potential RNA-binding domains, and other functionally relevant features. Given the encoder output *H*, the decoder generates the target RNA sequence 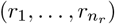 one token at a time. At each timestep *i*, the decoder conditions on, (i) the partially generated RNA sequence (*r*_1_, …, *r*_*i*−1_) via *self-attention*, (ii) the encoder output *H* via *cross-attention*:

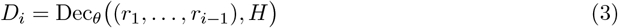

Here, *D*_*i*_ denotes the decoder’s hidden representation at timestep *i*. This vector *D*_*i*_ ∈ *R*^*d*^ captures contextual information from both the evolving RNA sequence and the encoded protein features. Finally, a linear layer *W*_vocab_ ∈ *R*^*d×*|*ν*|^ projects *D*_*i*_ onto the vocabulary space 𝒱 of RNA tokens, and a softmax function normalizes these scores to yield a probability distribution over all possible next tokens.

#### 4.3.2 Inference

At inference time, we produce a *de novo* RNA sequence 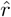 for a given protein *p* by sampling tokens sequentially from the output distribution:

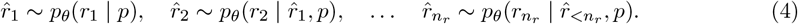

There are various techniques for sampling in language models [56, 57, 58]. For our work, we consider multiple approaches. One approach is greedy sampling, which selects the token with the highest probability at each time step. While this method often produces coherent sequences with high overall likelihood, it can be overly conservative and may lead to repetitive outputs. To mitigate this, we also use beam search. Beam search [56] maintains multiple candidate sequences by exploring the top *B* tokens (where *B* denotes the beam size) at each decoding step, eventually selecting the sequence with the highest overall probability. Although beam search generally improves sequence quality, it may still limit diversity. To introduce greater diversity, we incorporate stochastic sampling techniques such as top-*k* sampling and nucleus sampling (also known as top-*p* sampling). In top-*k* sampling [57], the token selection is restricted to the *k* most probable tokens, thereby reducing the likelihood of sampling low-probability tokens that could reduce sequence quality. Nucleus sampling, on the other hand, dynamically determines the candidate set based on a cumulative probability threshold *p*, ensuring that only the tokens contributing to the majority of the probability mass are considered. Both stochastic methods provide a balance between randomness and determinism, enabling the generation of RNA sequences that are diverse yet biologically plausible.

Recognizing that no single sampling strategy perfectly balances diversity and quality of the generated RNAs, we generate a pool of candidate RNA sequences using various sampling strategies and hyperparameter settings. We then apply scoring using predicted binding affinity, consistency, and minimum free energy (MFE). All of these metrics are normalized between zero and one, to score the generated sequences effectively. We normalize the MFE using min-max scaling 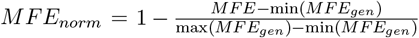 by this we ensure that lower MFE values are ranked higher. Binding affinity is derived from DeepCLIP model[34] and consistency measures the frequency of a given RNA sequence appearing across different sampling runs, normalized by dividing by the maximum occurrence observed 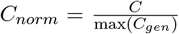. Higher consistency values indicate more robust sequence generation across sampling strategies.

Supplementary Figure 2 shows a comparative analysis of different sampling strategies using RNA pools generated for nine selected proteins. As shown in the figure, we evaluate several sampling strategies by adjusting hyper-parameters, including the top-k value and temperature, to determine the optimal combination that maximizes RNA quality scores. Among these sampling strategies, the top-k sampling method with *k* = 30 and temperature set to 1.5 consistently achieves the highest score. Given its superior performance, we employ this sampling strategy throughout all RNA generation and evaluation processes described in this study for RNAtranslator.

### 4.4 Experimental Setup

We trained RNAtranslator on two NVIDIA TITAN RTX GPUs using Fully Sharded Data Parallel (FSDP) to split the training across devices efficiently. we tokenize RNA and protein sequences using the Byte Pair Encoding (BPE) algorithm. A vocabulary size of 1,000 tokens was chosen to balance representation quality and computational efficiency. Each sequence is truncated or padded to a fixed length of 1,024 tokens to ensure uniformity during model training. We set a batch size of 8 and used gradient accumulation over 16 steps. We also applied weight decay with the Adam optimizer. The training carried on for 350,000 iterations over a month. Our hardware configuration used FP16 with optimization level O1 and a fixed seed of 42 for reproducibility. The model has a stack of 6 identical encoder-decoder layers with 12 attention heads. The total number of parameters is 41.4 million. In fine-tuning, we used three NVIDIA L40S GPUs, and increased the batch size to 15, conducting this phase for an additional 75, 000 iterations.

We trained GenerRNA on the same computational setup used for RNAtranslator. Specifically, we obtained a base GenerRNA model and fine-tuned it using RNA interactions from the CLIPdb database, employing 3, 917 interactions for RBM5 and over 1 million interactions for ELAVL1. We fine-tuned for 50, 000 iterations. For RNAGEN, we used the released pre-trained model and generated 500 sequences which are optimized to bind to RBM5 and ELAVL1 using the DeepBind model [14] for the corresponding protein.

### 4.5 Evaluation Metrics

Our primary objective is to ensure that the RNAs generated by RNAtranslator closely resemble natural RNAs in both structural properties and binding characteristics. To achieve this, we first define a set of properties that capture stability (e.g., minimum free energy, GC content) and binding specificity (e.g., affinity scores). We then compare the *distributions* of these properties with natural RNA sequences. To ensure a fair and comprehensive evaluation, we validate the generated RNAs by comparing their distribution. We use the DeepCLIP model[34] to quantify the RNA-protein binding affinity given a protein and RNA sequence. This model predicts the probability of binding based on features that correlate with known protein-binding motifs and RNA sequence patterns. For each target protein, we divide the RNA interaction dataset from CLIPdb into training, validation, and test sets with proportions of 80%, 10%, and 10%, respectively. We then train a DeepCLIP model using this partitioned data. For validation, we specifically select 500 RNAs from the test set as representatives of natural binding RNAs.

Furthermore, beyond sequence-based validation, we also analyze the designed RNAs at the structural level and simulate RNA-protein complexes using docking and moleculdar dynamics simulations. Since ground-truth structure for the designed RNAs are not available, we predict the structure using RhoFold+ [44]. Using HDOCKlite [45], we model the RNA-protein complex. HDOCKlite is a lightweight version of the HDOCK docking algorithm that focuses on RNA-protein interactions, incorporating a scoring function optimized for molecular docking [45]. Through these docking and MD analyses, we evaluated the interactions of the generated RNA sequences with their target proteins, assessing docking scores, potential complex structures, and the stability and dynamic properties of each RNA-protein interaction.

### 4.6 Molecular Dynamics Simulations

We use molecular dynamics simulation framework OpenMM [38] to assess structural dynamics of the designed RNAs. During the simulations, we employ a canonical ensemble (NVT) setup with a Langevin integrator at 300 K and a 2 fs time step. Each system undergoes energy minimization, followed by equilibration and a 10-million-step production run. Trajectories are recorded at regular intervals to track conformational changes over time. This standardized protocol is applied to all systems, including RNA–protein complexes and isolated RNAs. Before running simulations, we preprocess the structures with pdbfixer[46] to correct missing or nonstandard residues and remove unnecessary heterogens, then prepare using the AMBER14 force field, ensuring appropriate protonation states at pH 7.0.

To quantify binding affinity during simulations, we calculate the interaction energy (Δ*E*) using:

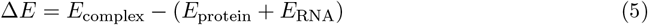

where *E*_complex_, *E*_protein_, and *E*_RNA_ represent energies of the RNA-protein complex, isolated protein, and isolated RNA, respectively. A more negative Δ*E* signifies stronger and more favorable interactions between the protein and RNA [59]. We further evaluate several structural and dynamic metrics to gain deeper insights into binding characteristics. Specifically, we quantify the number and stability of hydrogen bonds (H-bonds), as a higher number of stable H-bonds typically correlates with stronger binding interactions [60, 61, 62]. Hydrogen bonds are identified using a donor–acceptor distance cutoff of 4.5Å and a donor–hydrogen–acceptor angle cutoff of 150, following commonly accepted geometric criteria. Additionally, the number of atomic contacts is calculated between the two molecular components, where any pair of atoms within a distance of 0.45*nm*(4.5Å) is counted as a contact. A greater number of such close contacts generally corresponds to a larger and potentially more stable binding interface [62, 63].

We also computed the free energy landscape (FEL) based on molecular dynamics simulations by selecting a group of atoms in the protein and using the first frame of the simulation as a reference. For each subsequent frame, we calculate the root-mean-square deviation (RMSD) which measures how much the atomic positions differ from the reference and the radius of gyration (RG) which reflects the spread of the atoms around their center of mass. These two values are then used to construct a two-dimensional histogram that estimates the probability density of observing particular RMSD and RG combinations. To smooth out any noise, we apply a Gaussian filter to the histogram. The free energy landscape is calculated using the relation

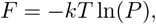

where *kT* represents the thermal energy (with *k* as the Boltzmann constant and *T* as the temperature) and *P* is the probability density from the smoothed histogram. By shifting the minimum free energy value to zero, we can clearly determine the depth of the landscape, which indicates the energy barrier between the most stable and less stable states. A larger depth indicates a higher energy barrier, meaning the complex is more stable [64, 65].

