## Supplementary Material for "RNAtranslator: Modeling protein-conditional RNA design as sequence-to-sequence natural language translation"

### Supplementary Material
### for
### RNAtranslator: Modeling protein-conditional RNA design as sequence-to-sequence natural language translation

#### 1 Supplementary Figures

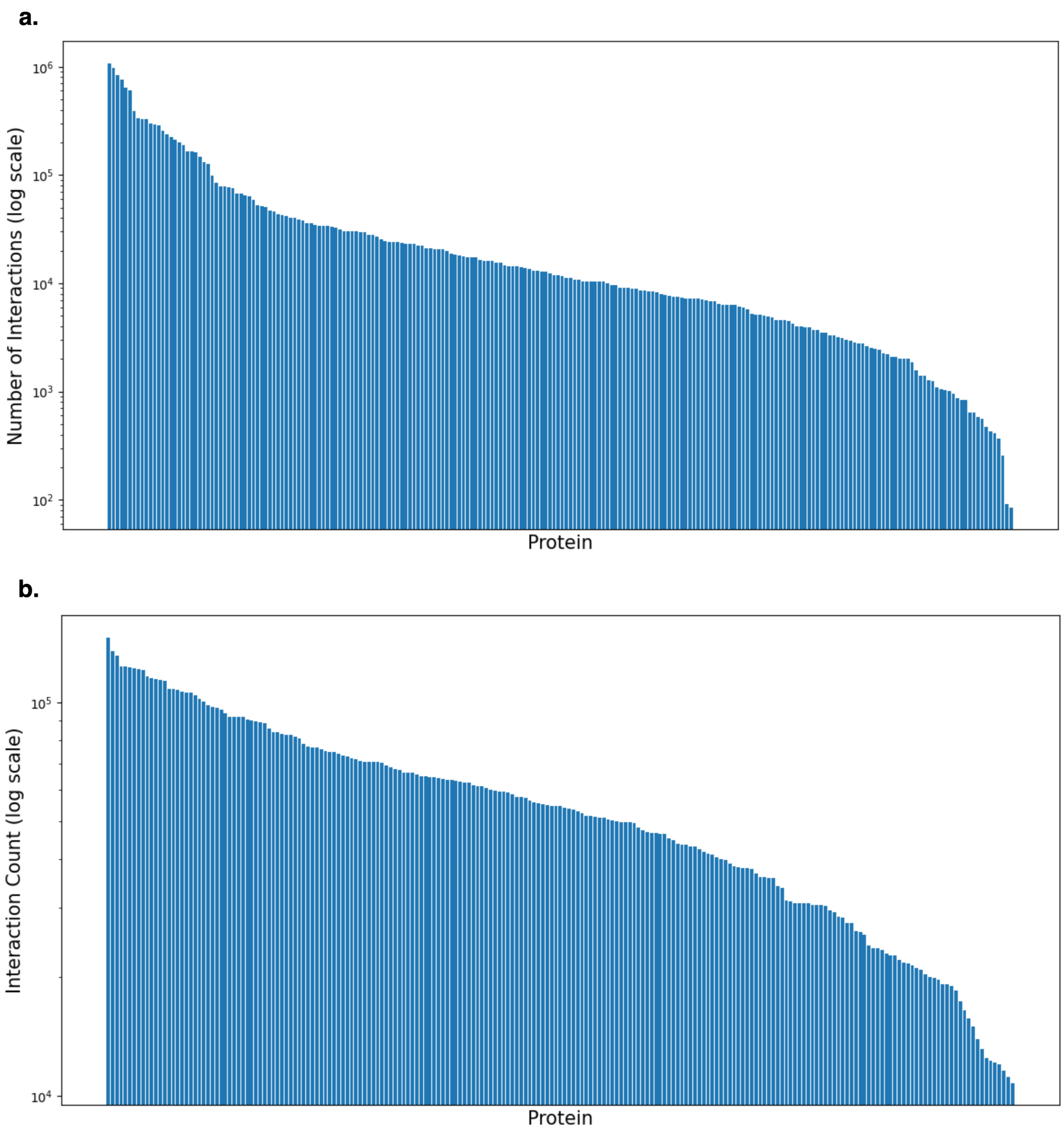


Supplementary Figure 1: **Analysis of CLIPdb dataset:** (a) Significant imbalance in interaction counts among proteins before preprocessing; (b) Balanced distribution of interactions achieved through oversampling and weighted sampling methods, maintaining the original diversity of RNAs.

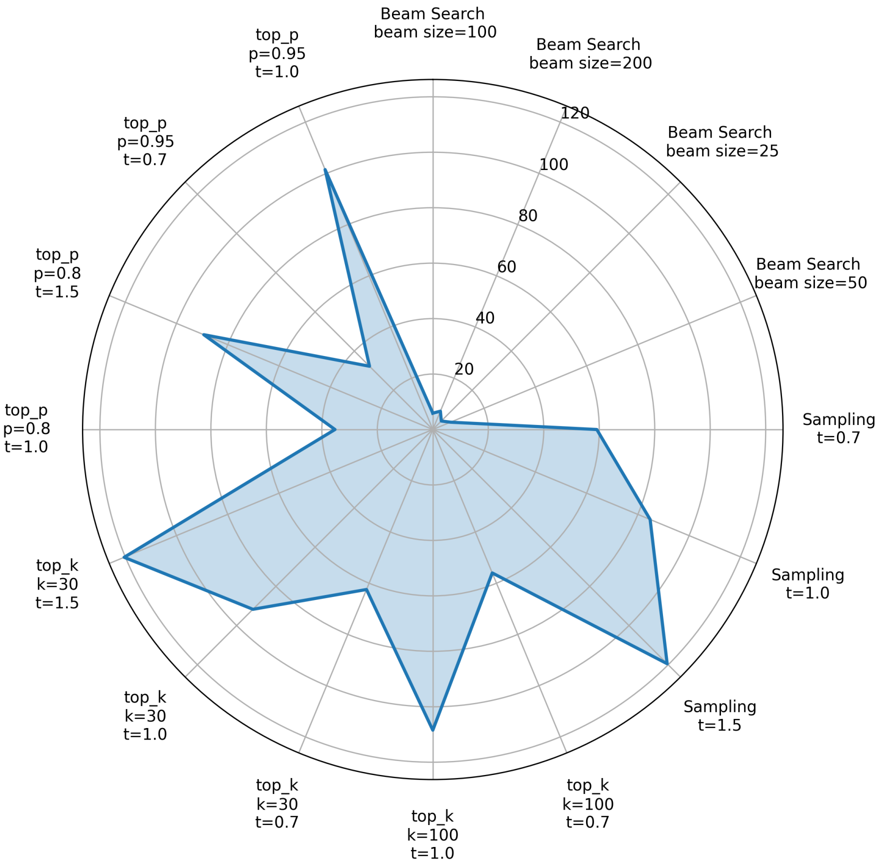


Supplementary Figure 2: **Comparison of RNA generation sampling strategies:** Scores for RNAs generated for nine proteins using different sampling approaches. The top-k sampling strategy with parameters $k = 30$ and $temperature = 1.5$ achieves the highest overall score.

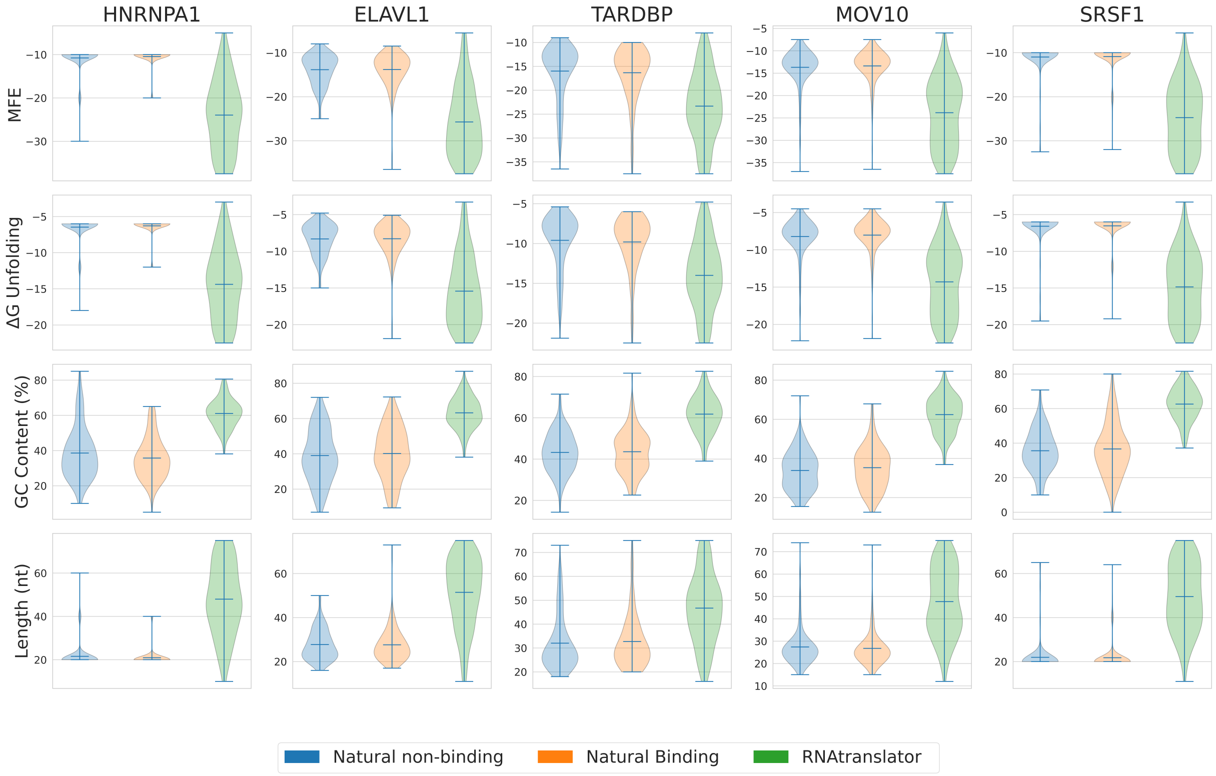


Supplementary Figure 3: **Stability analysis comparing RNAtranslator-generated and natural RNAs:** RNAtranslator-generated RNAs show similar Minimum Free Energy (MFE) and Ensemble Free Energy ($\Delta G$ ensemble) distributions to natural binding RNAs, indicating thermodynamic and structural stability. GC content analysis confirms structural robustness, and sequence length distribution highlights their broader range.
